# TFIIIC regulates SMC complex binding and 3D DNA contacts between tRNA genes

**DOI:** 10.64898/2026.04.27.721182

**Authors:** Daniel Obaji, Jun Kim, Yetunde Olagbegi, Anh-Thu Le, Sevinç Ercan

## Abstract

Transcription factor IIIC (TFIIIC) is a multi-subunit protein complex that recruits RNA polymerase III (Pol III) to the majority of its target genes. Evolutionarily conserved overlap between TFIIIC binding sites and the structural maintenance of chromosomes (SMC) complexes suggested a role for TFIIIC in SMC regulation and 3D organization of eukaryotic genomes; but the evidence has remained largely correlational due to the essential role of TFIIIC in recruiting RNA Pol III for transcription initiation. Here, we directly tested the function of TFIIIC in SMC complex regulation by using auxin inducible protein degradation in *C. elegans*. We performed Hi-C and ChIP-seq analyses upon acute depletion of TFTC-3, an essential TFIIIC subunit, and RPC-1, the catalytic subunit of RNA Pol III. Our results showed that TFIIIC is required for the binding of cohesin and condensin complexes to tRNA genes. TFIIIC is also required for increasing 3D contacts between distant tRNA genes located on the same or different chromosomes. Depletion of individual SMC complexes did not significantly reduce the tRNA gene contacts, suggesting redundancy or an independent mechanism of interphase genome organization mediating the 3D contacts between tRNA genes. Together, our study supports an RNA Pol III independent function for TFIIIC in regulating multiple SMC complexes and the 3D organization of tRNA genes.

## Introduction

Transcription factor IIIC (TFIIIC) recruits RNA polymerase III (Pol III) to a majority of its targets including transfer RNA (tRNA) and 5S ribosomal RNA (rRNA) genes (Ramsay & Vannini, 2018). At the tRNA gene promoters, TFIIIC recognizes DNA sequence motifs called the A-box and B-box located inside the transcribed region. At the 5S rRNA gene, TFIIIA binds to A- and C-box promoter elements and recruits TFIIIC. Recruitment of TFIIIC leads to the assembly of the transcription factor IIIB (TFIIIB) complex, including the TATA binding protein (TBP) (Ciganda & Williams, 2011; Kassavetis et al., 1990). RNA Pol III is recruited to a number of small nuclear (snRNA) genes including U6, signal recognition particle (SRP), and 7SK snRNA without TFIIIC (Murphy et al., 1987; Ramsay & Vannini, 2018; Schramm & Hernandez, 2002).

In addition to RNA Pol III transcribed genes, TFIIIC binds to distinct genomic sites without recruiting RNA Pol III (Hiraga et al., 2012; Moqtaderi et al., 2010; Moqtaderi & Struhl, 2004; Van Bortle et al., 2014). These sites, termed ‘extra TFIIIC’ (ETC) were found in multiple species including *S. cerevisiae* (Moqtaderi & Struhl, 2004)*, S. pombe* (Noma et al., 2006), *Caenorhabditis elegans* (Stutzman et al., 2020), *Drosophila melanogaster* (Van Bortle et al., 2014), mouse and humans (Moqtaderi et al., 2010; Yuen et al., 2017). In these species, ETCs are composed of diverged sequence elements partially resembling A and B boxes.

TFIIIC binding sequences were shown to be necessary and sufficient for the insulation function of specific tRNA genes in yeast (Noma et al., 2006; Raab et al., 2012; Valenzuela et al., 2009). In mammalian cells, insulation activity of tRNA genes depends on the genomic context (Donze & Kamakaka, 2001; Hamdani et al., 2019; Hong et al., 2024; Raab et al., 2012). This context may be linked to SMC complexes, as TFIIIC binding sites significantly overlap with that of multiple SMC complexes, and are enriched at the boundaries of topologically associating domains (TADs) (Dixon et al., 2012; Ferrari et al., 2020; Yuen et al., 2017).

While the overlap between TFIIIC and SMC complex binding led to the hypothesis that TFIIIC regulates 3D genome organization and SMC complex activity, the evidence remained largely correlational in part due to TFIIIC’s essential function in protein translation, as TFIIIC recruits RNA Pol III for transcription of tRNAs and 5S rRNA. Here, we directly tested this hypothesis by taking advantage of the auxin inducible system in *C. elegans* to acutely deplete TFIIIC and RNA Pol III, and elucidated an RNA Pol III-independent function for TFIIIC in regulating the binding of multiple types of SMC complexes.

We targeted TFTC-3 and RPC-1 for auxin inducible depletion in somatic cells of L3 larvae using *peft-3* driven TIR1. Incubating larvae on auxin-containing plates for one hour effectively reduced TFTC-3 but not RPC-1, providing a condition to study TFIIIC-dependent effects before reducing RNA Pol III transcription. ChIP-seq analyses of condensin subunits DPY-26 and DPY-27 and cohesin subunit SMC-3 showed that TFIIIC, but not RPC-1, is required for cohesin and condensin localization to tRNA genes. Hi-C analyses upon TFTC-3 and RPC-1 depletion suggested that both contribute to chromosomal separation of arm and centers, while TFIIIC is specifically required for increasing 3D contacts between distant tRNA genes. Together, our results establish that TFIIIC has a distinct role in controlling multiple SMC complexes and the 3D organization of tRNA genes.

## Methods

### Worm strains and growth

Worms were grown and maintained on NGM plates at 20-22°C with E.coli strains OP50 and/or HB101 bacteria. Synchronized L3 collections were made by bleaching gravid adults (0.5M NaOH and 1.2% sodium hypochlorite) and hatching embryos overnight without food in M9 (22 mM KH2PO4 monobasic, 42.3 mM Na2HPO4, 85.6 mM NaCl, 1mM MgSO4). The starved L1s were grown at 22°C for 22 hours. The motif scramble strain PHX6865 and degron-GFP tags were created by SUNY Biotech and sequence information is provided in Supplemental File 1. Depletion strains were generated by crossing to CA1200 to produce ERC94: syb5481[tftc-3::GGGGS::AID::emGFP] V; [eft3p::TIR1::mRuby::unc-54 3’UTR + Cbr-unc-119(+)] II and ERC100: syb6989 [rpc-1::GGGGS::AID::emGFP]IV; [eft3p::TIR1::mRuby::unc-54 3’UTR + Cbr-unc-119(+)] II.

### Auxin treatment

Auxin (indole-3-acetic-acid, Fisher 87-51-4) was resuspended in 100% ethanol to 400 mM. 1 mM auxin was added to NGM media before plates were poured. L3 larvae were washed three times with M9 and spread onto the auxin containing plate. After one hour, worms were collected using M9. For 22 hr collections, starved L1s were plated on auxin plates with food and grown for 22 hours at 22°C. For HiC and ChIP, worms were crosslinked in 2% formaldehyde for 30 minutes, followed by quenching in 125 mM glycine for 5 minutes, washing once with M9, two times with PBS+PMSF and protease inhibitors.

### DAPI and GFP imaging

Worms were washed at least twice with M9, crosslinked in 0.5% formaldehyde for 15 minutes, submerged in liquid nitrogen for 1 minute to freeze-crack, thawed in 37°C water bath and incubated on ice for 10 minutes. Worms were washed with PBS, then 70% ethanol, then 1 mL wash buffer (PBS-0.05% Tween-20), and resuspended in 1 mL wash buffer with 1 μL of 2 μg/mL DAPI and incubated in 37°C water bath for 30 minutes. After washing twice in PBS, worms were transferred to slides or agar pads for imaging. Images were acquired with a Zeiss Axio Imager A2 microscope and the AxioVision Rel.4.8 software.

### Worm sizing

The hatched and starved L1s were plated on plates with 1mM or without auxin, and with OP50 as food, and grown for 22 hours at 22°C. The worms were washed with M9 to remove bacteria, and resuspended in 40 mL of M9. Size measurements were obtained using the COPASTM FP-500 BioSorter, and exported to R for analysis. Time of flight, TOF, was used as a metric for axial length.

### ChIP-seq

ChIP-seq was performed as previously described (Albritton et al., 2017).100-200 μL of pelleted worms were washed once and dounce-homogenized with 30 strokes in FA buffer (50 mM HEPES/KOH pH 7.5, 1 mM EDTA, 1% Triton X-100, 0.1% sodium deoxycholate, 150 mM NaCl) with 0.1% sarkosyl and 1mM PMSF and 1X protease inhibitors Calbiochem cocktail I. Dounced worms were sonicated in Picoruptor to 200-800 bp. 1-2 mg of extract was used per ChIP and 5% was taken as input DNA. 5-10 μg of antibody per ChIP was collected by 20 μL of Protein A sepharose and washed 2 times with FA buffer 5 min, once with FA-1 M NaCl buffer 5 min, once with FA-500 mM NaCl buffer 10 min, once with TEL buffer (0.25 M LiCl, 1% NP-40, 1% sodium deoxycholate, 1mM EDTA, 10 mM Tris-HCl pH 8.0) 10 min, and 2 times with TE buffer 5 min. Immunoprecipitated chromatin was eluted from the beads by incubating with ChIP elution buffer (1% SDS, 250 mM NaCl, 10 mM Tris pH 8.0, 1 mM EDTA) at 65 °C for 15 minutes, incubated with 2 μL 10 mg/mL Proteinase K at 50 °C for 1 hour and reverse crosslinked overnight at 65 °C. DNA was purified with Qiagen Minelute PCR purification kit. Half of the ChIP DNA and 30 ng of input DNA were used for library preparation as previously described (Albritton et al., 2017).

### Hi-C

Hi-C was performed as previously described (Kim et al., 2022). Formaldehyde-crosslinked ∼100-150 μL worm pellet was resuspended in 40-60 μL of PBS and dripped into a mortar containing liquid nitrogen. The worms were ground with a pestle until a fine powder. Ground worms were crosslinked again with 2% formaldehyde in TC buffer following the Arima High Coverage HiC kit which uses four 4-base cutters: DpnII, HinfI, DdeI, and MseI or the Arima HiC kit which uses two 4-base cutters: DpnII and Hinfl. Arima’s protocol was followed for library preparation using KAPA Hyper Prep Kit.

### High-throughout sequencing

At least two biological replicates of ChIP-seq were sequenced using multiplexed single or paired-end 75 bp sequencing with Illumina NextSeq 500, NovaSeq, or Aviti. Replicate, read and mapping information for all sequenced samples are provided in Supplemental File 1. Two biological replicates of Hi-C were sequenced using paired-end 100 bp sequencing with the Illumina Novaseq 6000. All sequencing was performed by the GenCore facility at the New York University Center for Genomics and Systems Biology, New York, NY. The Illumina reads were basecalled using Picard IlluminaBasecallsToFastq version 2.23.8 (https://broadinstitute.github.io/picard/), with APPLY_EAMSS_FILTER set to false. Following basecalling, the reads were demultiplexed using Pheniqs version 2.1.0 (Galanti et al., 2021).

The entire process was executed using a custom nextflow pipeline, GENEFLOW (https://github.com/gencorefacility/GENEFLOW). Aviti reads were basecalled using Bases2Fastq version 2.1.0 (https://docs.elembio.io/docs/bases2fastq/). Following basecalling, the reads were demultiplexed using Pheniqs version 2.1.0 (Galanti et al., 2021). The entire process was executed using a custom nextflow pipeline, GENEFLOW.

### ChIP-seq data processing

Bowtie2 version 2.4.2 (Langmead & Salzberg, 2012) was used to align 75 bp single-end reads to WS220 or modified WS220 genome for the scramble mutant, with default parameters. Bam sorting and indexing was performed using samtools version 1.11 (Danecek et al., 2021). BamCompare tool in Deeptools version 3.5.0 (Ramírez et al., 2016) was used to normalize for the sequencing depth using CPM and create ChIP-Input coverage with a bin size of 10 bp and 200 bp read extension. Only reads with a minimum mapping quality of 20 were used, and mitochondrial DNA, PCR duplicates, and blacklisted regions were removed (Amemiya et al., 2019). The average coverage data was generated by averaging ChIP-Input enrichment scores per 10 bp bins across the genome. ChIP-seq peaks were identified using MACS2 version 2.1.1 (https://github.com/macs3-project/MACS). Peaks were called using the individual replicates with minimum false discovery rate of 0.05, and with the merged bam files with minimum false discovery rate of 0.01. Final peaks were generated using bedtools version 2.9.2 (Quinlan & Hall, 2010) to overlap the MACS2 peaks from the merged bam file with the peaks from the individual replicates such that a peak is considered final if it appears in the merged bam peaks and the majority of the replicates.

### Hi-C data processing

Hi-C data was mapped to the ce10 (WS220) reference genome using default parameters of the Juicer pipeline version 1.5.7 (Durand et al., 2016). The biological replicates were combined using juicer’s mega.sh script. The mapping statistics from the inter_30.txt output file are provided in the supplemental file 1. The inter_30.hic outputs were converted to cool format using the hicConvertFormat of HiCExplorer version 3.6 (Ramírez et al., 2018; Wolff et al., 2018, 2020) in two steps using the following parameters: 1) --inputFormat hic, --outputFormat cool, 2) -- inputFormat cool --outputFormat cool --load_raw_values. The cool file was balanced using cooler version 0.8.11 (Abdennur & Mirny, 2020) using the parameters: --max-iters 500, --mad-max 5, --ignore-diags 2. The balanced cool file was used for all downstream analysis. To get log-binned P(s) curves and the derivative, insulation, compartment strength, cooltools v0.7.0 (Open2C et al., 2022, 2024) (https://github.com/open2c/cooltools) was used. For off diagonal pileups, coolpuppy was used (Flyamer et al., 2020) (https://github.com/open2c/coolpuppy)

### Gene annotations

tRNA gene annotation was obtained from tRNA database (https://gtrnadb.org) and filtered to keep only high confidence genes and then lifted over to the ce10 genome using UCSC liftOver (Kuhn et al., 2013) with default parameters and using ce11ToCe10.over.chain chain file. Non-coding RNA genes were obtained from worm mine version WS289 and lifted over to the ce10 genome using UCSC liftOver.

### TFIIIC and RNA Pol III site classification

To define the three classes of genomic sites bound by TIIIIC and RNA Pol III, the TFTC-3 and RPC-1 final peaks from control strain were merged using bedtools version 2.9.2 (Quinlan & Hall, 2010). This resulted in a total of 2624 sites in L3 and 2066 in embryos. This annotation was used with deeptools mutliBigwigSummary (Ramírez et al., 2016) to get the average ChIP enrichment of TFTC-3 and RPC-1 over the sites. Bedtools intersect was used to get the overlap between the union annotation and high confidence TFTC-3, RPC-1 peaks, tRNA genes, and other ncRNA genes. For null comparison, a random list of sites with similar chromosomal distribution as the union of TFTC-3 and RPC-1 peaks was generated in R, then annotated tRNA and ncRNA genes were excluded from this to get a total of 2301 random sites. The enrichment over the random sites was used to set cutoffs for TFTC-3 and RPC-1 ChIP peaks that were considered as background. To obtain high confidence TFIIIC+,Pol III+ sites, union annotation sites that i) overlapped with both the high confidence TFTC-3 MACS peaks and RPC-1 peaks, and ii) had ChIP enrichment higher than the cutoff were selected. High confidence TFIIIC+,Pol III-sites were those final TFTC-3 peaks that were above the background cutoff and did not overlap with RPC-1 peaks or any gene annotation. The high confidence TFIIIC-,Pol III+ sites were the final RPC-1 peaks that were above the background cutoff and did not overlap with TFTC-3 or tRNA genes.

### Motif analysis

Bedtools (v2.29.2) getfasta was used to get the DNA sequence for the three annotation classes for both L3 and embryo. To get the motifs, MEME (v5.3.0) (Bailey et al., 2015) was used with the following options: -mod zoops, -nmotifs 5, -minw 6,, -objfun classic, - revcomp and -markov_order 0.

### Data and script availability

Sequence data are publicly available at the Gene Expression Omnibus (GEO) under GSE338224 for ChIP and GSE338222 for Hi-C. Scripts are available at https://github.com/ercanlab/2026_Obaji_TFIIIC_SMC.

## Results

TFIIIC is a six subunit protein complex organized into two domains: tauA and tauB which bind to the promoter elements box A and box B respectively (Girbig et al., 2022; Mylona et al., 2006; Vorländer et al., 2020). In *C. elegans* three out of the six TFIIIC subunits have been identified by sequence and structure conservation (**Supplemental Figure 1A**). Previous studies reported TFIIIB, TFIIIC and RNA Pol III binding in *C. elegans* embryos (Gerstein et al., 2010; Stutzman et al., 2020). Here, we used early-stage L3 larvae, composed mostly of postmitotic somatic cells and a few germ cells.

ChIP-seq profiles of TFTC-3, TFTC-5 (TFIIIC subunits), TBP-1 (TFIIIB subunit), and RPC-1 (RNA Pol III catalytic subunit) in L3 larvae were similar to those obtained in embryos (Gerstein et al., 2010; Stutzman et al., 2020) (**Supplemental Figure 1B**). We categorized the TFTC-3 and RPC-1 peaks into three groups: TFIIIC+, Pol III+ sites (n=690), TFIIIC-, Pol III+ sites (n=217), and TFIIIC+, Pol III-(n=224) (**Supplemental Figure 1C**). As expected, the majority of TFIIIC+, Pol III+ sites are tRNA and noncoding RNA genes (tRNA: 430, ncRNA: 97). The TFIIIC-, Pol III+ sites are TFIIIC independent RNA Pol III target genes and the TFIIIC+, Pol III-sites comprise the ETCs.

Although slightly fewer ETCs were identified in L3 larvae compared to embryos (224 vs 268), their genomic distribution showed the same distinct enrichment at specific chromosome arms that correlate with meiotic pairing centers (Phillips et al., 2009) (**Supplemental Figure 1D**).

DNA sequences corresponding to A and B boxes derived from the ETCs in L3 larvae were similar to those identified in embryos (**Supplemental Figure 1E**). Thus, TFIIIC binding at ETCs persists from the embryo to the L3 larval stage.

### Acute depletion of TFTC-3 and RPC-1 from somatic cells in L3 larvae

For auxin inducible depletion of TFIIIC, we targeted TFTC-3, a highly conserved and essential subunit that functions in the recruitment of TFIIIB (Giaever et al., 2002; Liao et al., 2003; Moir & Willis, 2004; Vorländer et al., 2020). For RNA Pol III, we targeted the catalytic subunit RPC-1. We endogenously tagged *tftc-3 and rpc-1* at the C terminus with degron-GFP and crossed to obtain homozygous strains expressing the auxin receptor TIR1 under the control of a soma-specific promoter *peft-3* (Zhang et al., 2015) (**Figure 1A**). In the absence of auxin, the resulting strains were superficially wild type with mild to no significant effect on viability (**Supplemental Figure 2A**).

**Figure 1:**
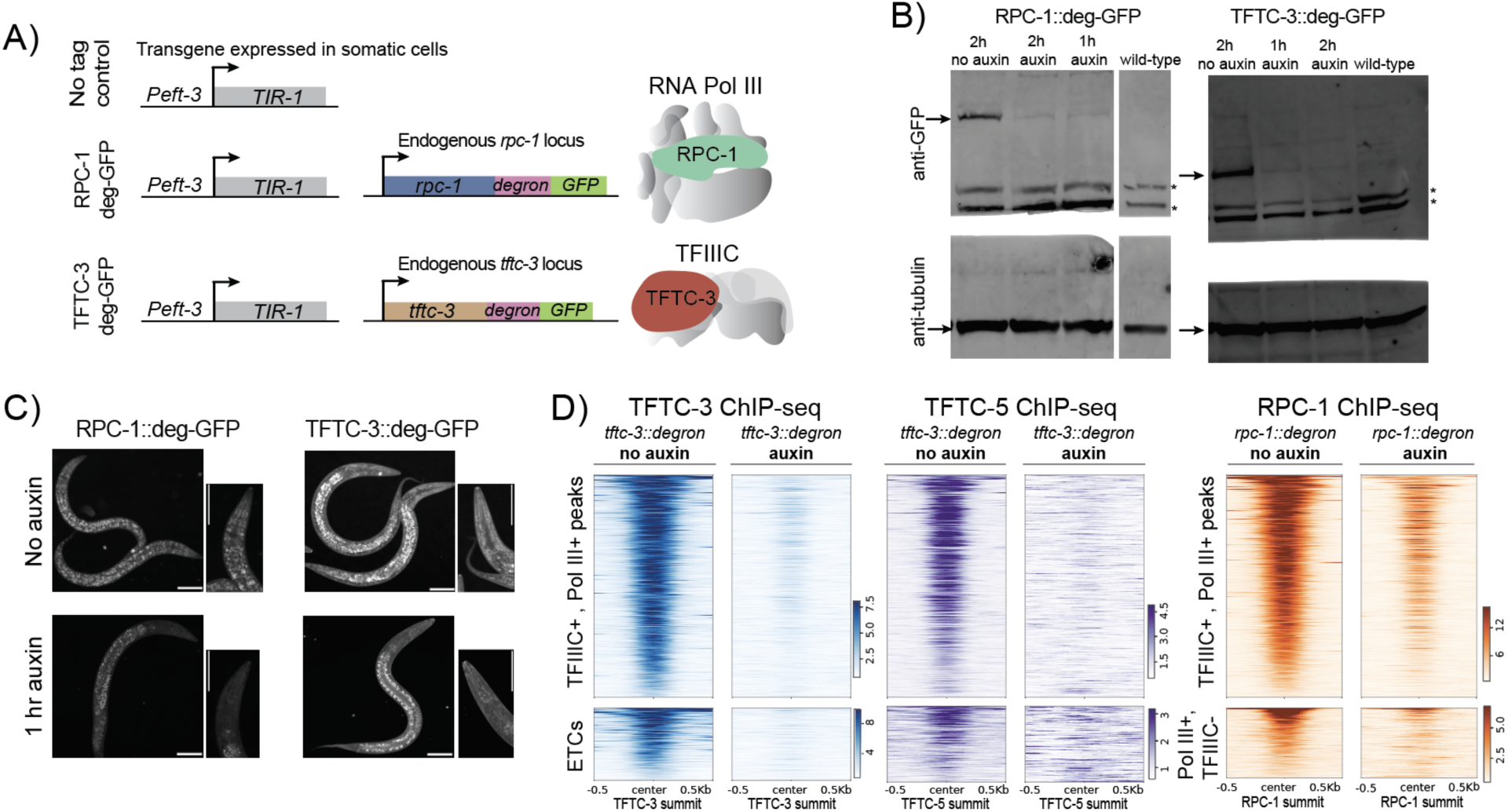
Auxin-inducible depletion of TFIIIC and RNA Pol III in somatic cells of *C. elegans.* (A) Schematic of the auxin inducible degron system used to deplete TFTC-3 and RPC-1. Endogenous *rpc-1* and *tftc-3* are tagged with degron and GFP at the C terminus, and TIR-1 is exogenously expressed under a somatic promoter *eft-3*. (B) Western blot analysis of extracts from L3 worms incubated on auxin containing plates for 1 hr. Anti-GFP antibody was used to detect the tagged protein, and anti-tubulin antibody was used as loading control. (C) Images show depletion of GFP-tagged proteins following 1 hour auxin treatment. GFP signal is lost from somatic cells but retained in germline cells following auxin treatment in both *tftc-3* and *rpc-1* tagged strains. Scale bar is 50 μm.(D) Heatmap showing ChIP enrichment scores across the different groups of TFIIIC and RNA Pol III peaks. Data from no auxin control is shown. TFTC-3 and TFTC-5 ChIP enrichment are plotted upon 1 hr auxin depletion of TFTC-3. RPC-1 ChIP enrichment is plotted upon 1 hr depletion of RPC-1.

Western blot analysis of TFTC-3 and RPC-1 demonstrated inducible depletion as early as one hour of incubating worms on auxin-containing plates (**Figure 1B**). Soma-specific depletion was validated by the reduction of the GFP signal across the body except the few mitotic germ cells in early L3 larvae (**Figure 1C**). We also confirmed the genome-wide depletion of TFTC-3 and RPC-1 by ChIP-seq analysis, and validated that TFTC-3 degradation leads to reduced binding of another TFIIIC subunit TFTC-5 (**Figure 1D**).

### TFTC-3 and RPC-1 depletion impede development

Interestingly, 1 hr depletion of TFTC-3 did not cause a significant reduction in RPC-1 binding (**Figure 2A**). Longer depletion of TFTC-3 by incubating L1 larvae on auxin-containing plates for 22 hours reduced RPC-1 binding, validating that TFIIIC is required for RNA Pol III transcription (**Figure 2B**). The prolonged depletion of TFTC-3 and RPC-1 did not kill, but significantly retarded growth (**Figure 2C**). RPC-1 depleted larvae remained small with two germ cells undivided after 22 hrs while TFTC-3 depleted larvae displayed some germ cell division (**Figure 2D**). It is likely that tRNAs, with half lives in the order of several days (Nwagwu & Nana, 1980; Karnahl & Wasternack, 1992; Berg & Brandl, 2021), support the viability of TFTC-3 and RPC-1 depleted larvae.

**Figure 2:**
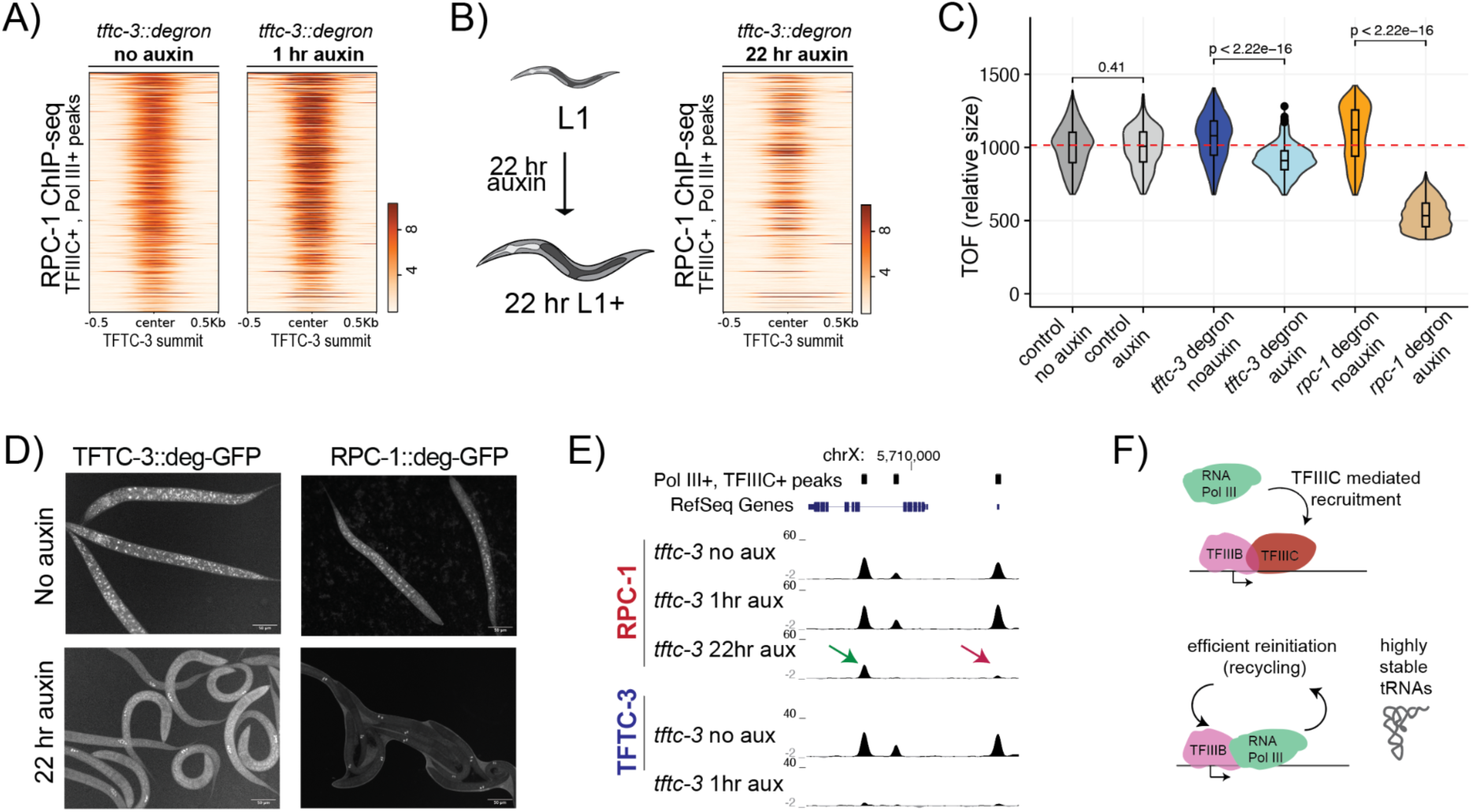
RNA Pol III binding persists after 1 hr depletion of TFIIIC but reduce after 22 hr. (A) Heatmap showing RPC-1 ChIP enrichment across RPC-1 peaks upon 1 hr knockdown of TFTC-3 knockdown in early L3 larvae. (B) 22 hr depletion of TFTC-3 was accomplished by growing synchronized L1 larvae on auxin containing plates. RPC-1 ChIP signal reduced upon 22 hr TFTC-3 depletion. (C) Violin/boxplots of time of flight (TOF), which is a metric for worm length, is plotted for L1s grown for 22 hours on plates containing no auxin and 1mM auxin. (D) GFP signal in L1 larave grown on auxin plates mark the germ cells, as *peft3*::TIR1 drive GFP-degron tagged TFTC-3 and RPC-1 degradation in the soma. (E) UCSC genome browser view including two sites with more (green) and less (red) persistence of RNA Pol III following TFIIIC depletion. (F) RNA Pol III recycling (re-initiation of transcription without TFIIIC), combined with high stability of tRNAs likely support the viability and development of the 22 hr TFTC-3 depleted larvae.

### Acute depletion of TFTC-3 provides support for recycling of RNA Pol III *in vivo*

The persistence of RPC-1 binding upon one hour TFTC-3 degradation is consistent with the idea that previously bound RNA Pol III reinitiate transcription, termed “recycling” (Dieci & Sentenac, 1996; Ferrari et al., 2004; Cabart et al., 2008; Arimbasseri et al., 2014). RPC-1 persistence varied between genes (**Figure 2E**).To analyze differences between RNA Pol III promoter types, we grouped genes based on their promoter sequence (**Supplemental Figure 2B**). We reasoned that efficient recycling would reduce the correlation between the change in RPC-1 binding and TFTC-3 depletion, which was 0.36 for tRNA, 0.38 for ncRNA, and 0.52 for snoRNAs (**Supplemental Figure 2C**). Gene length increases the reliance on TFIIIC for initiation (Ferrari et al., 2004). RPC-1 retention negatively correlated with the length of snoRNA genes (**Supplemental Figure 2D**), thus providing further evidence that RNA Pol III is recycled *in vivo* (**Figure 2F**).

### TFIIIC and RNA Pol III contribute to 3D separation of chromosome arms and center

*C. elegan*s autosomes are roughly separated into two arms and a center (Gerstein et al., 2010). The center domains are relatively gene rich and the arms are enriched in heterochromatin and lamina contacts (Ikegami et al., 2010). The X chromosome contains a single arm on the left, while the rest of the chromosome is located towards the interior of the nucleus (Snyder et al., 2016). Depletion of TFTC-3 and RPC-1 had minimal impact on average contact frequency (**Supplemental Figure 3A**) and the organization of the genome into chromosomes (**Figure 3A**).

**Figure 3:**
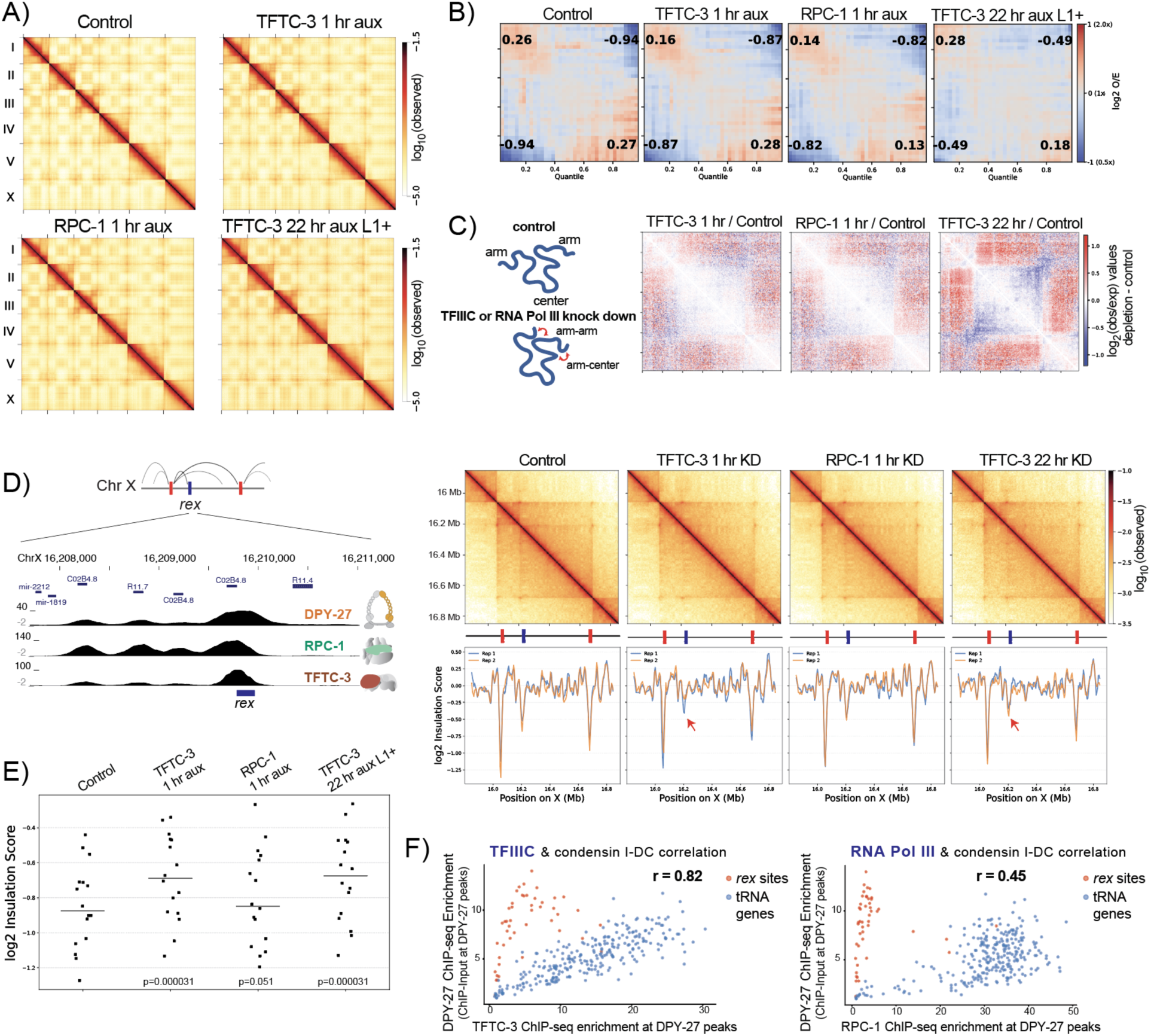
TFIIIC and RNA Pol III knockdown effects on 3D organization of chromosomes and X chromosomal TADs. (A) HiC matrix of the genome at 50kb resolution in no tag control, TFTC-3 1 hour and 22 hour knockdown (L1+ refers to L1s incubated on auxin plates with food for 22 hr), and RPC-1 1 hour knockdown. (B) PCA analysis of compartmentalization. Observed/expected values of 2×2 bins at the top-left corner (B-B), bottom-right corner (A-A), top-right and bottom-left corners (A-B) of the saddle plots. (C) Change in Hi-C contacts with normalized to no tag control on chromosome III. (D) Condensin I-DC recruitment elements on the X (*rex*) form TAD boundaries. ChIP-seq and Hi-C data surrounding a *rex* site near a cluster of four tRNA genes. (E) Average insulation scores at the 17 strong *rex* sites are plotted for each condition. Wilcoxon rank sum test test was performed against no tag control. (F) Scatter plot of DPY-27 versus TFTC-3 and RPC-1 ChIP enrichment over tRNA genes (blue) and *rex* sites (red).

However, separation of chromosome arms and centers slightly reduced upon TFTC-3 or RPC-1 depletion, as quantified by arm/center compartmentalization scores (**Figure 3B**) and relative to no tag control (**Figure 3C**). Smaller chromosomes I, II and III, which have a more defined arm and center separation showed a stronger effect, while chromosomes IV, V and X displayed more complicated domain structures (**Supplemental figure 3B**). Notably, the effect upon TFTC-3 depletion was observed for all chromosome arms regardless of ETC enrichment (**Supplemental Figure 3B**), suggesting that the TFIIIC function in isolating arm and center organization of the chromosomes is not specific to ETCs.

We reasoned that the longer depletion of TFTC-3, which also reduces RPC-1 binding, should combine the effects of TFIIIC and RNA Pol III. Indeed, there was a stronger increase between arm and center contacts upon 22 hr of TFTC-3 depletion (**Figure 3C**). Together, these results suggest that both TFIIIC and RNA Pol III contribute to the 3D organization of chromosomes.

### TFIIIC effect on condensin I-DC mediated TAD boundaries on the X chromosomes

In *C. elegans*, X chromosomes contain loop-anchored topologically associating domains (TADs), which are formed by a condensin I variant through mechanisms akin to that of mammalian cohesin (Anderson et al., 2019; Crane et al., 2015; Kim et al., 2022). Condensin I^DC^ (hereafter I-DC) shares four subunits with condensin I and contains a paralog of SMC-4 called DPY-27 (Aharonoff et al., 2024; Csankovszki, 2009). Condensin I-DC is targeted specifically to the X chromosomes by a set of ***r***ecruitment ***e***lements on the ***X*** (*rex*) (Albritton et al., 2017; Fuda et al., 2022). A subset of the *rex* sites function as TAD boundaries by blocking condensin I-DC loop extrusion (Anderson et al., 2019; Rowley et al., 2020; Kim et al., 2022).

At a *rex* site near a cluster of four tRNA genes, we noticed higher TFTC-3 binding compared to RPC-1 (**Figure 3D**). Insulation across this *rex* site slightly decreased upon TFTC-3 depletion. Across all seventeen strong *rex* sites, the reduction of insulation was statistically significant for TFTC-3 but not RPC-1 (**Figure 3E**). This reduction does not reflect a general effect on chromatin insulation across the genome, because acute depletion of TFTC-3 decreased and increased insulation of 264 and 348 1kb-bins, respectively (FDR<0.01). Across the X chromosome, DPY-27 ChIP-seq signal correlated more with TFTC-3 (r = 0.8211) compared to RPC-1 (0.4478) (**Figure 3F)**. Furthermore, unlike the chromosome arm and center separation, 22 hr depletion of TFTC-3, which combines the effect of TFIIIC and RNA Pol III, did not further decrease insulation (**Figure 3E**), thus the observed reduction in insulation across the TAD boundaries is TFIIIC specific.

### TFIIIC regulates condensin I-DC binding to tRNA genes

In previous work, we observed that condensin I, I-DC and condensin II localize to tRNA genes (Kranz et al., 2013). To test if this binding required tRNA promoter sequences box A and box B, we scrambled the box A and B motifs at the X chromosomal tRNA gene F56C3.t1 (**Figure 4A**). Mutation of the tRNA promoter sequences eliminated ChIP-seq signal for TFIIIC and RNA Pol III and diminished the binding of condensin I-DC subunits DPY-26 and DPY-27 (**Figure 4A, 4B)**.

**Figure 4:**
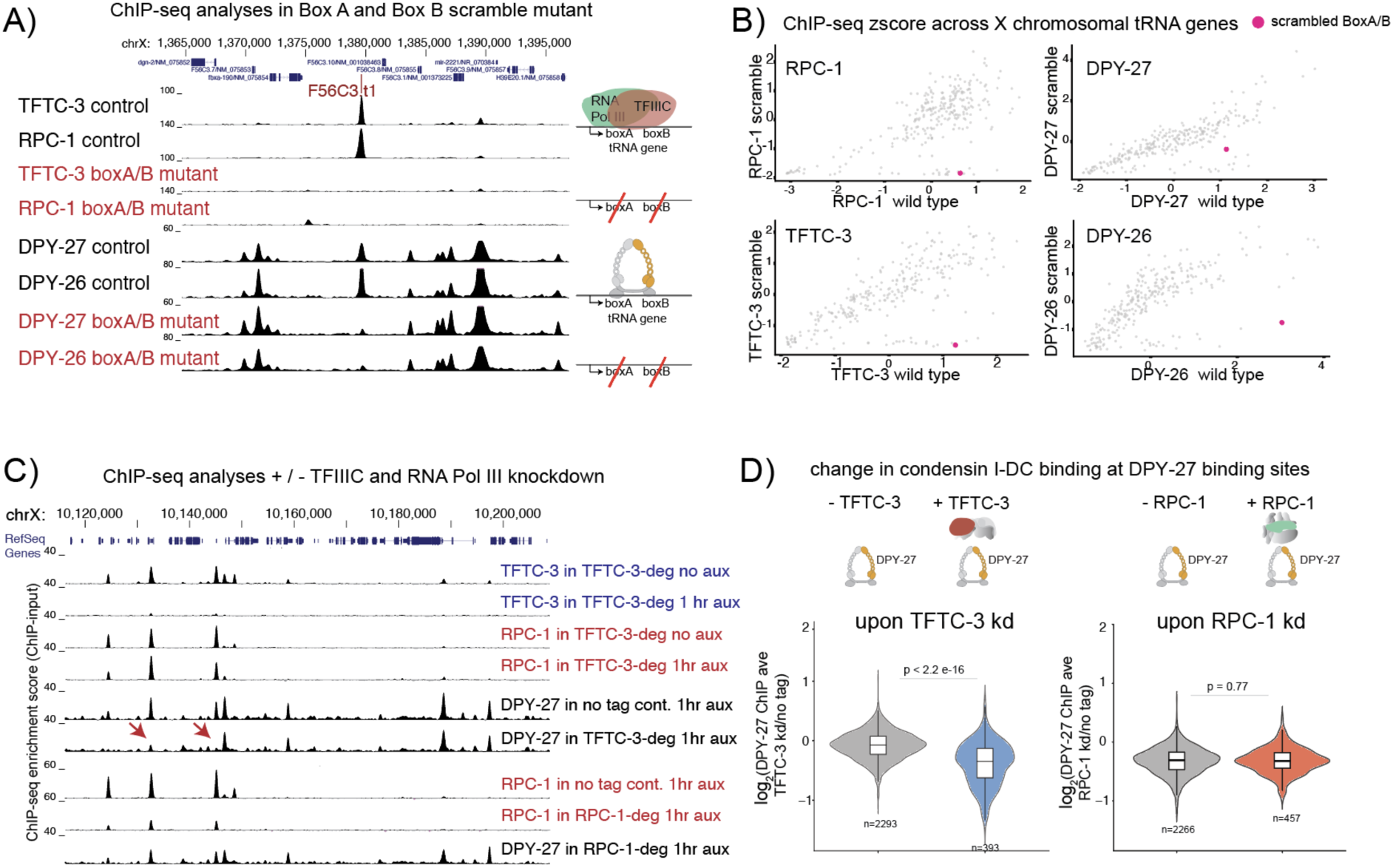
TFIIIC is required for condensin I-DC binding to X chromosomal tRNA genes. (A) UCSC genome browser view of the region surrounding the mutated A and B boxes at the tRNA gene F56C3.t1. TFTC-3, RPC-1, DPY-26 and DPY-27 ChIP-seq profiles in control and mutant are shown. (B) Scatter plot of average ChIP-seq enrichment over tRNA genes on the X chromosome. The mutated tRNA gene is marked in pink. The x axis is the average ChIP binding for the indicated protein in the wild-type and the y axis is the same in the mutant strain. (C) UCSC genome browser view of a representative region showing ChIP-seq profiles of TFTC-3, RPC-1, and DPY-27 in control, TFTC-3 depletion and RPC-1 depletion conditions. Red arrows highlight two TFIIIC-associated sites where DPY-27 binding is specifically diminished following TFTC-3 depletion. (D) Violin and boxplots of the log2 fold change in DPY-27 binding following TFTC-3 depletion (left) and RPC-1 depletion (right). DPY-27 peaks were categorized based on overlap with TFTC-3 (left) and RPC-1 (right) peaks in the no tag control.

Next, we asked if depletion of TFIIIC or RNA Pol III reduced condensin I-DC. Importantly, only a subset of condensin I-DC binding sites are associated with tRNA genes, thus DPY-27 peaks that do not overlap with TFIIIC or RNA Pol III serve as an internal control to assess the effect of TFTC-3 and RPC-1 depletion. Upon depletion of TFTC-3, condensin I-DC binding is reduced specifically at the DPY-27 peaks that overlap with TFTC-3, as shown for a representative region **(Figure 4C**) and quantified across the X chromosome **(Figure 4D)**. Together, these results show that TFIIIC specifically mediates condensin I-DC binding to the X chromosomal tRNA genes.

### TFIIIC mediates cohesin and condensin I binding to tRNA genes

To test if TFIIIC also regulates other SMC complexes, we performed ChIP-seq analysis of the cohesin subunit SMC-3. Strikingly, SMC-3 binding was also reduced only at the TFIIIC bound cohesin sites upon TFTC-3 depletion, shown across a representative region (**Figure 5A**) and quantified across the whole genome (**Figure 5B)**. In previous work, we showed that cohesin is preferentially loaded at/near active enhancers for RNA Polymerase II (Kim et al., 2025a). In line with TFTC-3 specifically regulating cohesin at TFIIIC sites, Hi-C fountains did not significantly change upon 1 hr depletion of TFTC-3 (**Supplemental Figure 4B**).

**Figure 5:**
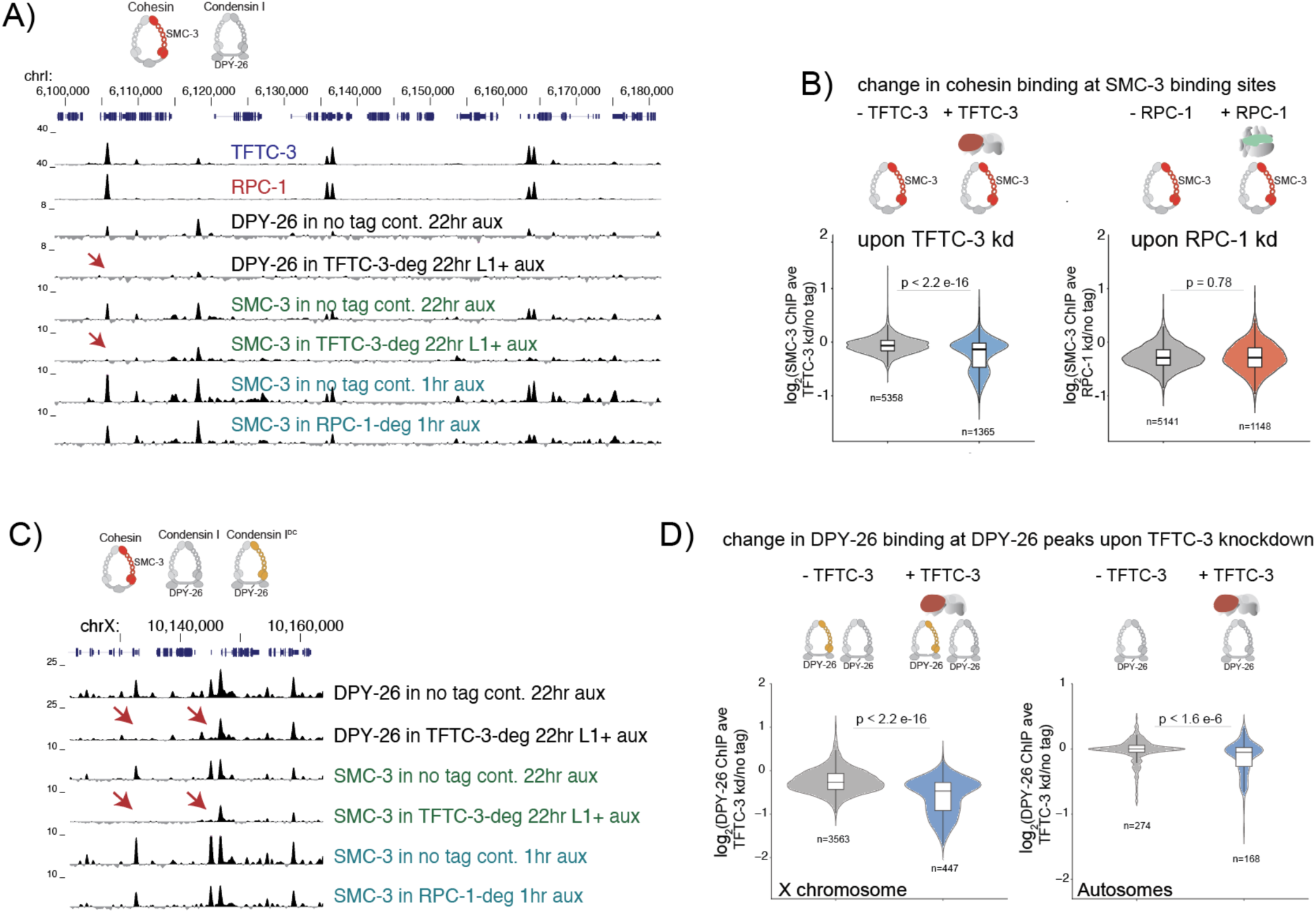
TFIIIC regulates binding of cohesin and condensin I at tRNA genes. (A) UCSC browser view of a representative autosomal region. ChIP-seq profiles of cohesin subunit SMC-3 and condensin I/I-DC subunit DPY-26 are shown in control and depletion conditions. Red arrows highlight an example site where SMC binding is reduced specifically in the TFTC-3 depletion. (B) Violin and boxplots of the log2 fold change in SMC-3 binding following TFTC-3 (left) and RPC-1 (right) depletion. SMC-3 peaks were categorized based on overlap with TFTC-3 (left) and RPC-1 (right) peaks in no tag control. (C) UCSC browser view of a part of the X chromosomal representative region shown for DPY-27 in Fig 4F. ChIP-seq profiles of cohesin subunit SMC-3 and condensin I/I-DC subunit DPY-26 are shown. Cartoons of the SMC protein complexes on top highlight the presence of the extra loop extruder condensin I-DC on the X chromosome. (D) Violin and boxplots of the log2 fold change in DPY-26 binding following TFTC-3 depletion for X chromosome peaks (left) and autosome peaks (right). All p-values are from Wilcoxon test.

We also analyzed condensin I, targeting DPY-26, a subunit of both I and I-DC (**Figure 5C)**. As expected, DPY-26 binding to autosomes was weak in largely interphase cells of L3 larvae. Nevertheless, there was a significant reduction in DPY-26 upon TFTC-3 depletion, specifically at DPY-26 peaks that overlap with TFIIIC. Together, these results highlight that TFIIIC controls the localization of multiple SMC complexes, including condensin I, I-DC and cohesin.

### TFIIIC promotes 3D DNA contacts between distant tRNA genes

Proteins that are barriers to SMC-mediated DNA loop extrusion localize SMC complexes and form distant loops observed in Hi-C matrices (Banigan & Mirny, 2020). Such loops exist between the *rex* sites on the X chromosomes (Anderson et al., 2019; Kim et al., 2022). Browsing the Hi-C interaction matrices did not reveal comparable loops between TFIIIC sites. However, pairwise contact frequency between tRNA genes was higher than expected (**Figure 6A**) and significantly reduced upon depletion of TFTC-3 but not RPC-1 (**Figure 6B**). Unlike tRNA genes, RNA Pol III sites that do not contain TFIIIC or a set of randomized sites did not show enrichment of 3D contacts (**Figure 6C**). Thus, TFIIIC is specifically required for promoting 3D contacts between the TFIIIC bound tRNA genes.

**Figure 6:**
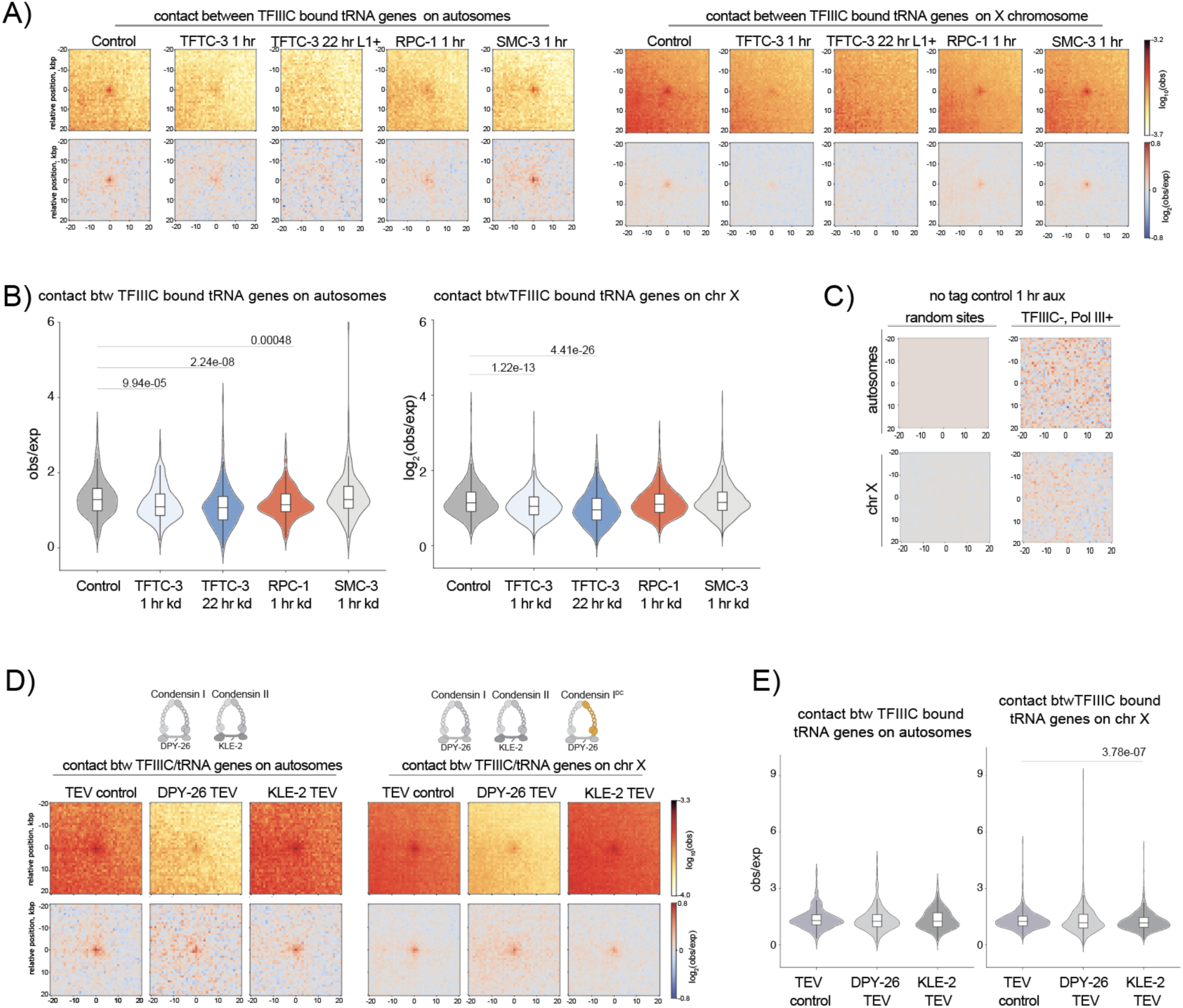
TFIIIC mediates long range 3D contacts between tRNA genes. (A) Off-diagonal pileup of HiC contacts between TFIIIC bound tRNA genes on autosomes and X chromosome in different experimental conditions. 1kb resolution matrices and pair-wise interactions that are distant 100-500 kb are shown. (B) Quantification of *cis* contacts between tRNA genes shown in panel A. For each valid pair of tRNA gene, the average observed/expected enrichment in a 3 by 3 square centered on the pair locus is plotted in violin/boxplots. A paired wilcoxon test with bonferroni correction was performed and the p values for significant comparisons are displayed. (C) Off-diagonal pileup of HiC contacts from randomized sites and TFIIIC-, Pol III+ sites on autosomes and the X chromosome. (D) Off-diagonal pileups as in panel A for TEV control, DPY-26 and KLE-2 cleave conditions from (Das et al., 2024). The SMC protein complexes with the relevant subunits for autosomes and the X chromosomes are displayed on top. (E) Violin/boxplot of the quantification of the contacts in panel D.

We next asked if TFIIIC-dependent contacts are mediated by a specific SMC complex. For cohesin, we used Hi-C data from 1 hr auxin-induced depletion of SMC-3 in L3 larvae (Kim et al., 2025b). For condensin I and II, we used data using TEV cleavage of DPY-26 and KLE-2 (Das et al., 2024). Depletion of cohesin, condensin I/I-DC and condensin II individually did not significantly reduce the 3D contacts between tRNA genes, as observed in Hi-C pileups (**Figure 6A** and **D**), and pairwise contact frequencies (**Figure 6B** and **E**).

## Discussion

In this study, we directly tested a long-standing hypothesis that TFIIIC regulates 3D genome organization independent of RNA Pol III transcription in multicellular eukaryotes. Below we discuss our results in the context of previous studies and speculate on the mechanisms by which TFIIIC may regulate SMC complex binding and mediate 3D contacts between tRNA genes.

### TFIIIC and RNA Pol III function in 3D organization of C. elegans chromosomes

*C. elegans* chromosomes are roughly organized into multi megabase arm and center domains. Depletion of TFTC-3 or RPC-1 did not eliminate but weakened this large-scale compartmentalization (Figure 3). The increased interactions between chromosome arms and centers upon TFIIIC or RNA Pol III depletion was similar to that seen following depletion of CEC-4, a nuclear envelope protein that mediates anchoring of chromosome arms to the nuclear lamina (Bian et al., 2020). In *C. elegans*, RNA Pol III transcribed loci associate with the nuclear pores and both TFTC-3 and RPC-1 immunoprecipitate the pore subunits NPP-13 and NPP-16 (Ikegami & Lieb, 2013). In addition, TFIIIC bound ETCs are enriched within lamin-associated protein LEM-2 domains (Stutzman et al., 2020). It is possible that TFIIIC and RNA Pol III reduce chromosomal contacts by mediating interaction with the nuclear lamina and pores.

### TFIIIC mediates localization of SMC complexes to tRNA genes independent of RNA Pol III

Here, we showed that TFIIIC regulates the binding of cohesin and condensin. In many organisms, including *D. melanogaster* and *C. elegans*, TFIIIC colocalize with condensins (D’Ambrosio et al., 2008; Kranz et al., 2013; Yuen et al., 2017; Yuen & Gerton, 2018). In mouse and human cells, tRNA genes and ETCs are enriched at the boundaries of TADs (Dixon et al., 2012; Yuen et al., 2017) and TFIIIC co-immunoprecipitates with the CCCTC-binding factor (CTCF) (Ferrari et al., 2020). In *S. pombe*, the TFIIIB subunit TBP physically interacts with condensin and localizes it to tRNA genes (Iwasaki et al., 2015). In human cell extracts, condensin II co-immunoprecipitates with TFIIIC subunits (Yuen et al., 2017). Our work in the context of these studies support an evolutionary conserved function for the general initiation factors, rather than RNA Polymerase III itself, in regulating SMC complexes.

TFIIIC may control the localization of cohesin and condensin by stalling DNA loop extrusion. A prediction of this hypothesis is that distant TFIIIC sites should interact at higher frequency than expected. Indeed, we observed an enrichment of pair-wise contacts between TFIIIC bound tRNA genes (**Figure 6A**). A second prediction of stalling is that the loop extruders would mediate an indirect interaction between TFIIIC and the sites where the loop extruder is stalled. Indeed, we observed TFIIIC at condensin I-DC *rex* sites that do not contain box A or B motifs (**Figure 3F**). A third prediction of regulating loop extruders is that TFIIIC should increase insulation of DNA contacts. We did observe a small but statistically significant reduction in insulation at *rex* sites upon TFTC-3 depletion (**Figure 3E**). Yet this effect was not genome-wide thus, TFIIIC must be a weak barrier to loop extrusion, and its contribution is observed only at sites with high extruder activity.

### Does loop extrusion alone explain TFIIIC mediating 3D contacts between tRNA genes?

Compared to DNA contacts that occur in *cis* (intra chromosomal), loop extrusion models have more difficulty explaining *trans* (inter chromosomal) interactions mediated by SMC complexes. An example of this is in *S. cerevisiae*, where tRNA genes on different chromosomes cluster in a condensin dependent manner (Iwasaki et al., 2010, 2015; Noma et al., 2006; Paul et al., 2018, 2019). In *C. elegans*, we find three observations that are difficult to explain if we assume TFIIIC functions *only* by regulating SMC loop extrusion.

First, although subtle, Hi-C contacts are enriched between TFIIIC binding sites on different chromosomes (**Figure 7A**). Second, while redundancy may explain the maintenance of tRNA gene contacts upon depleting individual SMC complexes (**Figure 6**), TEV cleavage of DPY-26, a subunit of both condensin I and I-DC, argues that an additional mechanism is required: If the extra loop extruder (I-DC) on the X chromosomes mediated tRNA gene clustering, then DPY-26 TEV cleavage should have reduced the Hi-C contacts on the X chromosomes more, compared to those on the autosomes, which is not the case (**Figure 6D**). The third observation is a lack of 3D contacts between tRNA genes in embryos, where SMC complexes are expressed and functional (**Figure 7B**).

**Figure 7:**
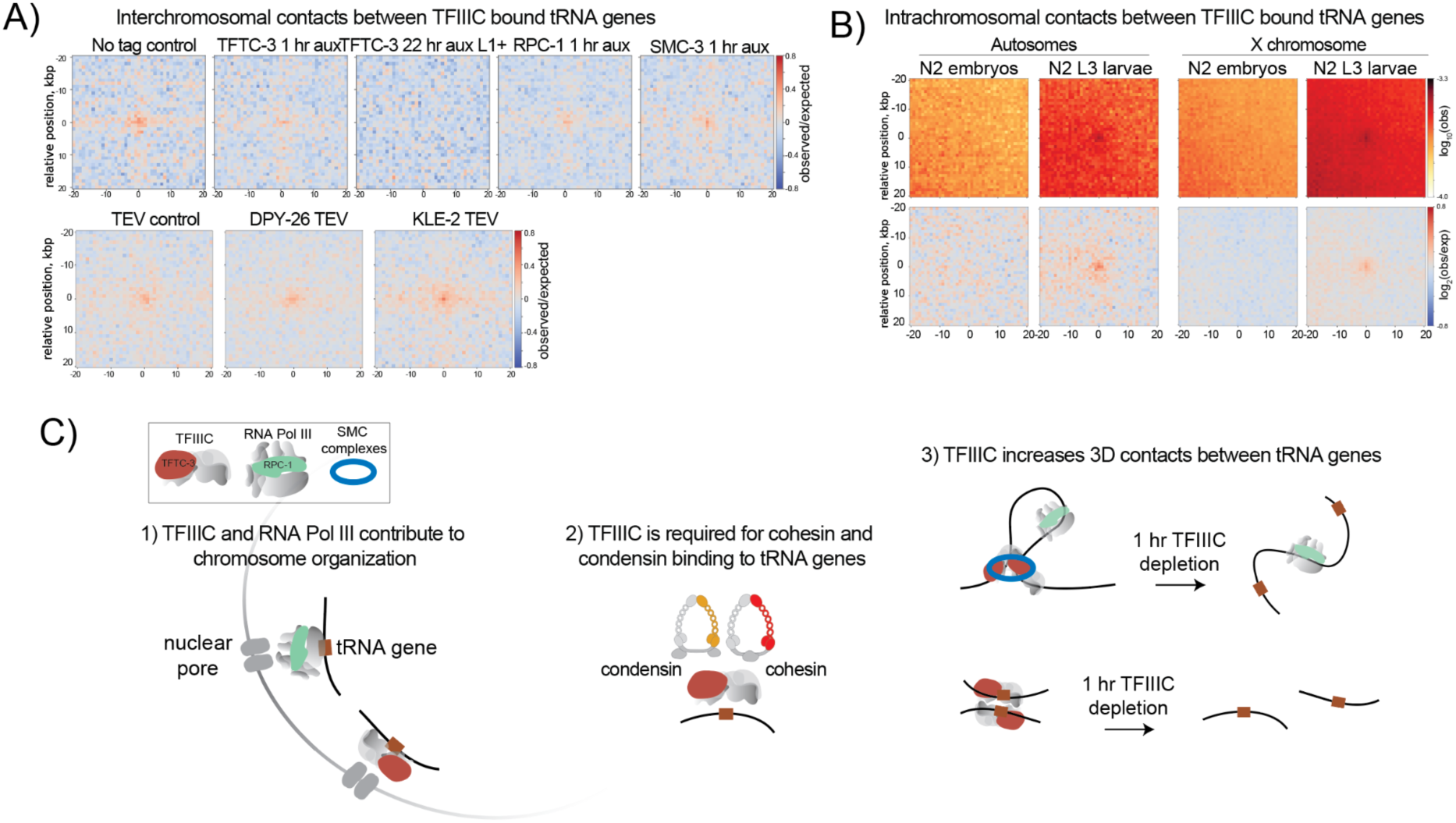
A model for the mechanisms of TFIIIC mediated 3D contacts between tRNA genes. (A) Pile up of observed/expected contacts between TFIIIC bound tRNA genes located on different chromosomes. (B) HiC contacts between TFIIIC bound tRNA genes on autosomes and X chromosome in wild-type (N2) embryos and L3 larvae. (C) TFIIIC and RNA Pol III may reduce 3D chromosomal contacts, possibly by mediating interaction with pores. TFIIIC, but not RNA Pol III, is required for the binding of cohesin and condensin to tRNA genes. TFIIIC-mediated contacts between tRNA genes are infrequent and likely heterogeneous within the population of cells. TFIIIC may be a weak barrier to loop extrusion by cohesin and condensin, thus localizing SMC complexes at a fraction of tRNA genes in a given cell. Presence of tRNA gene contacts between different chromosomes in larvae and the lack of contacts in embryos suggest that an SMC-independent, but TFIIIC-dependent mechanism of interphase genome organization contributes to the 3D contacts between tRNA genes.

Therefore, another mechanism must contribute to the 3D contacts between tRNA genes. It has been proposed that interphase chromatin has an intrinsic propensity to self-organize and bring active elements together (Mirny et al., 2019; Schooley et al., 2026). This self-organization and loop extruders may initially bring TFIIIC sites together and protein-protein interactions between TFIIIC or associated proteins may maintain the contacts (**Figure 7C**). The lack of tRNA gene clustering in embryos is consistent with this model, as the cells in embryos undergo continuing cell divisions that resets the chromatin landscape.

## Acknowledgements

DO, JK and SE, and research in this manuscript were supported by NIGMS of the National Institutes of Health under award number R35 GM130311. YO and AL were supported in part by NYU dean’s undergraduate research funds. We thank Gencore at the NYU Center for Genomics and Systems Biology for sequencing and raw data processing and acknowledge the Zegar Family Foundation for their generous support.

---

**Supplemental Figure 1:**
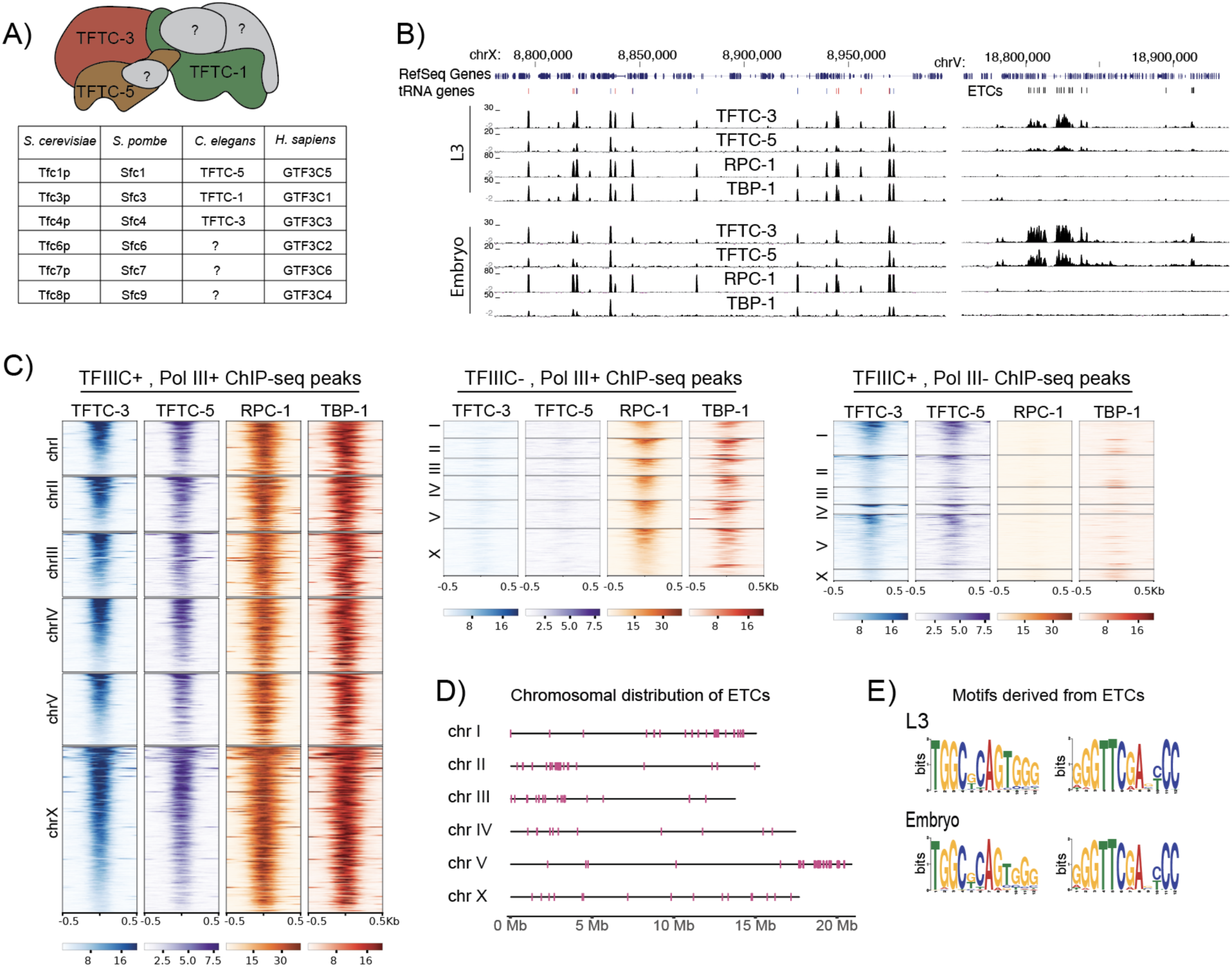
TFIIIC and RNA Pol III have shared and distinct binding sites in the *C. elegans* genome (A) The identified TFIIIC complex subunits in *C. elegans* are shown along with the yeast and human orthologs. (B) ChIP-seq enrichment profiles of TFTC-3, TFTC-5 (TFIIIC subunits), RPC-1 (RNA Pol III subunit), TBP-1 (TFIIIB subunit) in larval stage L3 (this study), and embryos (modENCODE) are shown at representative regions on the X chromosome and chromosome V. y axis is ChIP-Input normalized coverage and the x axis are chosen to show tRNA genes bound by TFIIIC and RNA Pol III, and ETCs bound only by TFIIIC. (C) Heatmap showing ChIP-seq enrichment of TFTC-3, TFTC-5, RPC-1, TBP-1 over a 1kb region centered on peak summits grouped as TFIIIC+, Pol III+ peaks, TFIIIC+, Pol III-peaks, and TFIIIC-, Pol III+ peaks. TFIIIC+, Pol III+ sites are overrepresented on the X chromosomes due to ∼45% of tRNA genes residing on the X in *C. elegans*. (D) Per chromosome distribution of ETC sites in larval stage L3 of *C. elegans*. (E) Sequence motifs generated using MEME with ETCs in the L3 versus embryos.

**Supplemental Figure 2:**
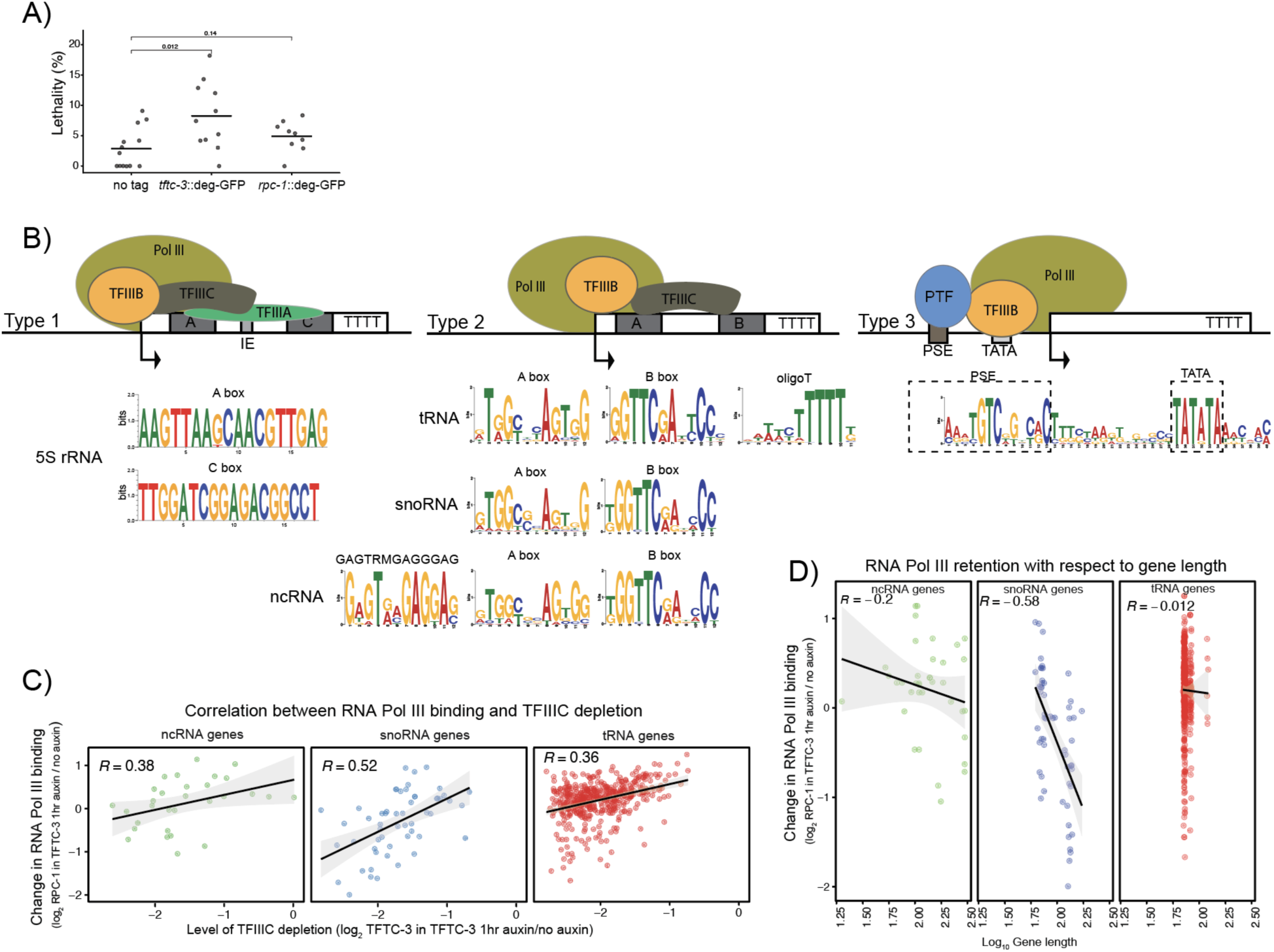
(A) The TFTC-3 and RPC-1 degron tagged strains were superficially wild type in the absence of auxin. Embryonic lethality was calculated as the percentage of unhatched embryos. Gravid adults were allowed to lay ∼20-40 embryos on 35 mm plates for 1-3 hours, and removed. The number of embryos were counted and the plates were kept at 20℃ for 24 hours to count the number of hatched worms. (B) *C. elegans* 5S rRNA, a type 1 gene, is transcribed from a cluster of repeats on the right arm of chromosome V (Nelson & Honda, 1985), a blacklisted region for ChIP-seq analysis (Amemiya et al., 2019). Pol III+, TFIIIC+ peaks overlap with the type 2 promoter genes encoding tRNA (442), snoRNA (55), other noncoding RNA (ncRNA) genes (30), or were unannotated (163 peaks). In addition to the previously recognized A and B box sequences, the ncRNA peaks bound by both RNA Pol III and TFIIIC revealed a new motif with the consensus GAGTRMGAGGAG (10/30 of sites, E value 4.5e-002) that is upstream of and present with both the A- and B-box motifs (9/10 times). Pol III+, TFIIIC-peaks overlapped with ncRNA genes (46) or were previously unannotated (166 peaks). Prior to this study, *C. elegans* was known to have a PSE at Pol II transcribed snRNAs and the Pol III transcribed U6 snRNA (Thomas et al., 1990). Our analysis shows the presence of a TATA box upstream of the TSS, thus *C. elegans* U6 snRNA genes likely utilize a strategy similar to vertebrates to specify Pol III transcription (Dergai & Hernandez, 2019). (C) Change in RPC-1 ChIP enrichment with respect to change in TFTC-3 depletion. The R values are pearson coefficients. (D) Change in RPC-1 ChIP enrichment upon 1 hr of TFTC-3 depletion with respect to gene length.

**Supplemental Figure 3:**
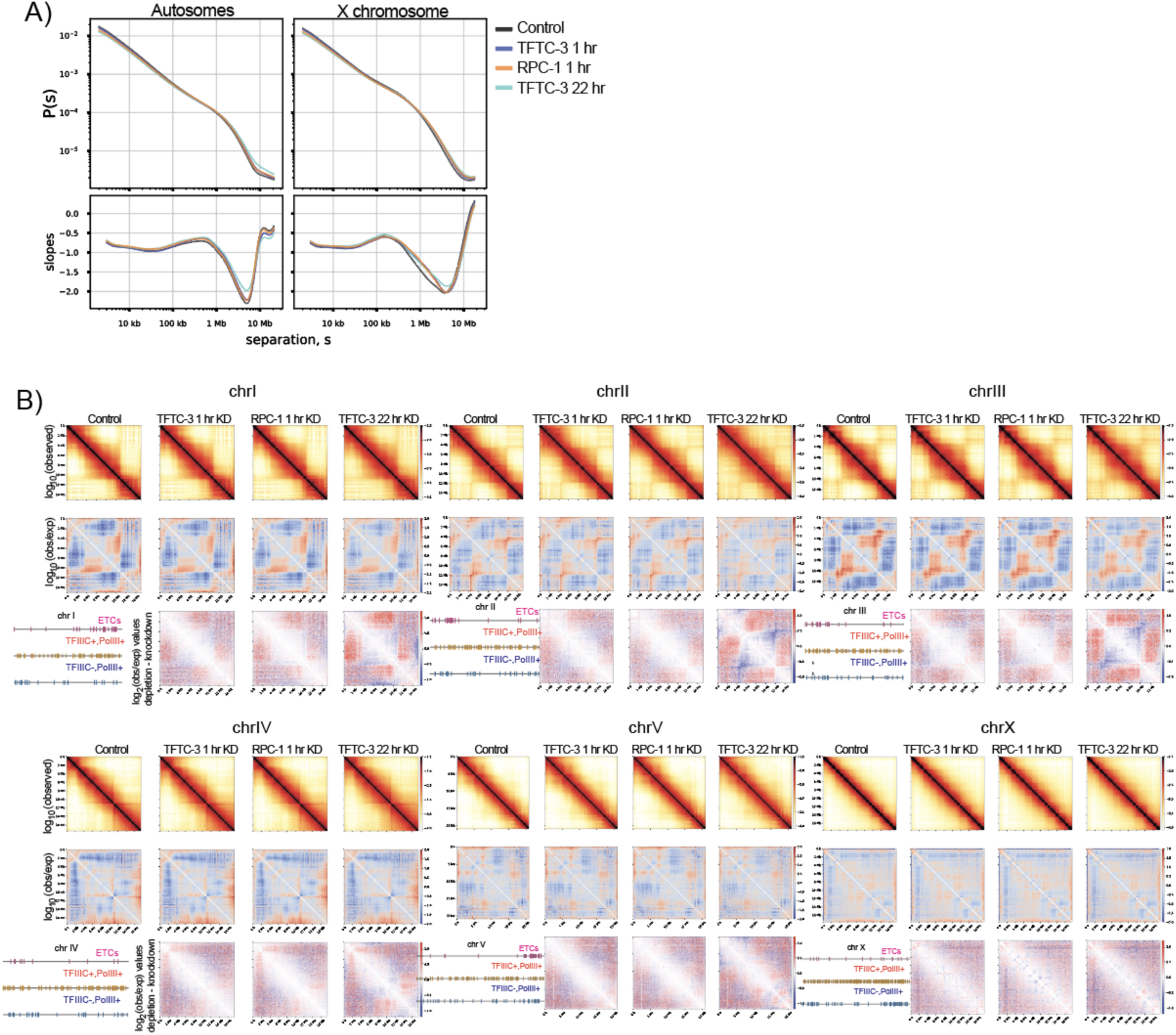
TFIIIC and RNA Pol III knockdown increases contacts between chromosome arms and centers. (A) Distance decay curves and slopes split by autosomes and chromosome X for control and depletion conditions. (B) Change in Hi-C contact matrix across a representative short autosome. The distribution of peak categories along the chromosome is shown on the upper left. Cartoon interpretation of the increase in contacts between the chromosome arms and center following TFIIIC and RNA Pol III knock down.

**Supplemental Figure 4:**
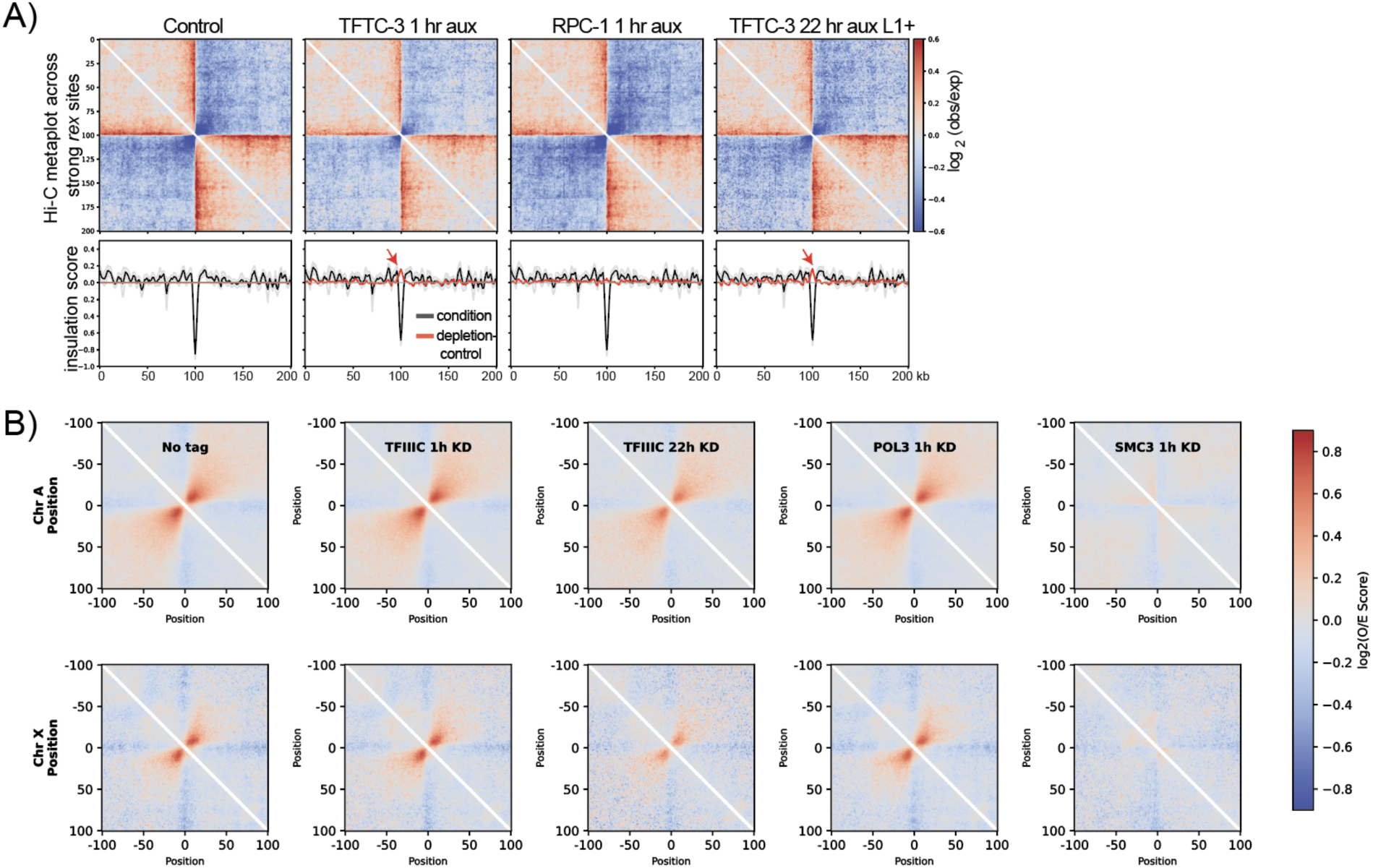
Effect of TFIIIC depletion on X chromosomal TADs and Hi-C fountains (A) On-diagonal pileup of observed/expected Hi-C matrix centered at the X chromosomal TAD boundaries (strong *rex* sites) and average insulation scores using 20kb window and 5kb resolution matrices. Black lines are the insulation scores for each condition labeled above, while the red line is the insulation score in the experimental condition minus the control. Red arrows point to increased insulation scores at the *rex* sites in the TFIIIC knockdown conditions. (B) On-diagonal pileup of observed/expected matrix centered on cohesin-mediated Hi-C fountain origins split into autosomes and chromosome X (top). The reduced contact frequency forming cohesin-mediated fountains upon 22 hr TFIIIC depletion may be due to indirect effects of prolonged depletion.

